# CHANGE-seq-BE enables simultaneously sensitive and unbiased *in vitro* profiling of base editor genome-wide activity

**DOI:** 10.1101/2024.03.28.586621

**Authors:** Cicera R. Lazzarotto, Varun Katta, Yichao Li, Elizabeth Urbina, GaHyun Lee, Shengdar Q. Tsai

**Affiliations:** Department of Hematology, St. Jude Children’s Research Hospital, Memphis, TN, USA; Beam Therapeutics, Cambridge, MA, USA

## Abstract

Base editors (**BE**) enable programmable conversion of nucleotides in genomic DNA without double-stranded breaks and have substantial promise to become new transformative genome editing medicines. Sensitive and unbiased detection of base editor off-target effects is important for identifying safety risks unique to base editors and translation to human therapeutics, as well as accurate use in life sciences research. However, current methods for understanding the global activities of base editors have limitations in terms of sensitivity or bias. Here we present CHANGE-seq-BE, a novel method to directly assess the off-target profile of base editors that is simultaneously sensitive and unbiased. CHANGE-seq-BE is based on the principle of selective sequencing of adenine base editor modified genomic DNA *in vitro*, and provides an accessible, rapid, and comprehensive method for identifying genome-wide off-target mutations of base editors.

## Main

Base editors^1,2^ are a transformative genome editing technology that can precise install single base alterations into cellular genomic DNA without requirements for DNA double strand breaks (DSBs), exogenous DNA donor templates, or reliance on cellular homology-directed repair pathways (HDR). For example, adenine base editors (ABEs)^2^ are capable of reverting G•C to A•T mutations, which represent the most common pathogenic SNPs reported in the ClinVar database (∼48%)^3^, by precisely installing targeted A•T to G•C point mutations in the DNA, highlighting the enormous potential of this class of genome editors for therapeutic application.

Base editors can bind and modify off-target genomic loci with sequence homology to the target protospacer, similar to other genome editors^4–6^. As base editors rapidly advance towards clinical application, defining guide RNA (gRNA) dependent off-target activity of adenine base editors in a sensitive and unbiased way is important for ensuring the safety of these new therapeutic approaches^7^. We consider as unbiased, methods that do not pre-select candidate off-target sequences for experimental analysis.

Current methods for profiling the genome-wide activity of base editors^8–10^ have limitations. Digenome-seq^8^ and EndoV-seq^9^ are based on whole genome sequencing of base editor treated genomic DNA *in vitro*, followed by processing with *E. coli* Endonuclease V^11^ to generate DSBs that can be detected by high-throughput sequencing. As these methods do not enrich for base editor modified genomic DNA, they require hundreds of millions of sequencing reads, reducing their sensitivity, scalability and increasing costs. Critically, the validation rate of these methods for identifying *bona fide* cellular off-target base editing mutations is relatively low^8,9^. Another recent method, ONE-seq^10^, is based on biased computational pre-selection of candidate off-target sites and, therefore, cannot identify genome-wide off-target activity beyond those sites selected *a priori*. Additionally, sensitive and unbiased approaches can offer the unique advantage of detecting unintended on-target genome editor off-target activity caused by ‘unknown unknowns’ such as the presence of contaminant gRNAs.

In earlier studies, we and others have applied indirect methods to identify base editor off-target activity^12–16^, relying on the simplifying, but not necessarily true, assumption that base editor off-targets are a subset of nuclease off-targets. For example, to experimentally nominate candidate off-target sites for an ABE therapeutic strategy to treat sickle cell disease (**SCD**), we performed CIRCLE-seq^17,18^ (an earlier method we developed for selective sequencing of Cas9 cleaved genomic DNA) using Cas9 nuclease to nominate candidate off-target sites. We then measured editing activity at CIRCLE-seq-Cas9 nominated off-target sites in a base editing context in CD34^+^ HSPCs treated with ABE8e-NRCH:*HBB*^S^-gRNA using multiplex targeted sequencing. We identified 54 Cas9-nuclease nominated off-target sites with unintended off-target base editing activity as high as 82%.

Although Cas-dependent adenine base editing off-targets often occurs at Cas9 nuclease off-targets^8,9,19^, it is known that productive base editing requires additional criteria that affect the off-target profile, such as the presence of adenine within the editing window, nucleotide sequence context, and R-loop accessibility by deaminases, that are not satisfied for all Cas-nuclease-dependent off-target sites^20^. Thus, limitations of our initial nuclease-centric approach are that base editors may have different off-target activity than cognate Cas9 nucleases, highlighting an unmet need for a simultaneously sensitive and unbiased method for specifically defining base editors genome-wide off-target activity. Critically, unbiased methods to characterize gene therapy products that utilize base editing to gain regulatory approval for clinical trials will need to be performed with the relevant base editor and not a similar nuclease.

Previously, we developed CHANGE-seq^21^, a sensitive, unbiased, and high-throughput method to identify Cas9 nuclease off-target activity *in vitro*. CHANGE-seq leverages a Tn5 tagmentation-based workflow for efficient generation of circularized genomic DNA libraries, coupled with stringent selection of circular DNA molecules using a cocktail of exonucleases, thereby, generating a population of circular DNA molecules with minimal free DNA ends. Covalently-closed circular DNA molecules are treated with CRISPR-Cas9 ribonucleoprotein complexes, and nuclease-cleaved DNA fragments are then ligated to adapters and sequenced, enabling the selective sequencing of Cas9 cleaved DNA molecules.

Here we present CHANGE-seq-BE, a method for both sensitive and unbiased, genome-wide discovery of guide-dependent base editor on- and off-target sites, adapted from CHANGE-seq^21^. After extensive molecular biology optimization, we overcame substantial technical challenges in enriching for editor-modified DNA reported by us and others^10,21^. CHANGE-seq-BE now enables the selective and efficient sequencing of base editor modified genomic DNA.

We focused our initial studies on adenine base editors because they exhibit lower guide-independent off-target activity and applied CHANGE-seq-BE to define the gRNA dependent genome-wide off-target activity of ABE8e in therapeutically relevant target sites in primary human cells. Strikingly, we found that CHANGE-seq-BE sensitively identifies not only off-target activity but also unintended on-target activity of contaminating synthetic gRNAs, highlighting one of the advantages of using unbiased sensitive methods for genome-wide assessment of genome editing activity.

## Results

### Development of a sensitive and unbiased biochemical method to measure base editor genome-wide activity

Motivated by a critical need for better, more direct methods to characterize the genome-wide off-target activity of base editors, we sought to develop a method that would be simultaneously sensitive and unbiased, and straightforward in practice. We reasoned that CHANGE-seq could be adapted for base editors by treating purified, circularized genomic DNA with base editor ribonucleoprotein (**RNP**) complexes and selectively sequencing base editor modified genomic DNA (**Fig. 1a**). First, linear genomic DNA could be circularized using our previously described CHANGE-seq Tn5-tagmentation mediated approach^21^. Second, enzymatically purified genomic DNA circles could be treated with ABE RNP complexes to nick the target DNA strand and deaminate adenine bases to inosine (within the base editing window) on the non-target strand at on- and off-target sites. Third, nicked and inosine-containing genomic DNA circles could be further processed with Endonuclease V^11^ which cleaves DNA adjacent to inosines to produce linear DNA with 5’ staggered ends. Finally, the base editor modified, endonuclease V processed linear DNA could be end-repaired and ligated to sequencing adapters for high-throughput sequencing.

**Figure 1.**
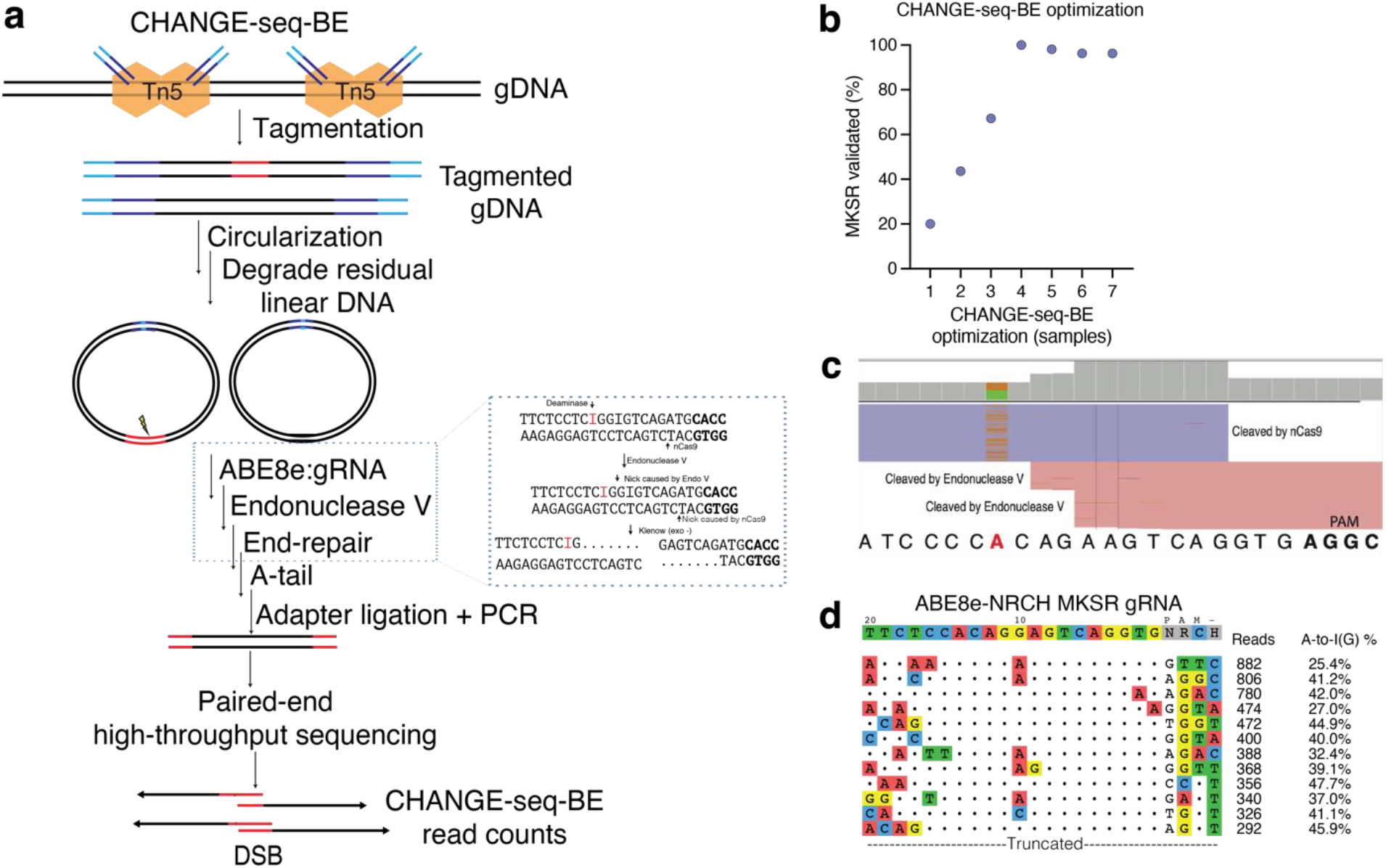
Development and optimization of CHANGE-seq-BE, a sensitive and unbiased biochemical method to detect base editor genome-wide activity. **a,** Schematic of CHANGE-seq-BE workflow. Purified genomic DNA (gDNA) is tagmented with custom Tn5 transposomes containing circularization adapter DNA. Tagmented DNA is prepared for circularization and circularized by intramolecular ligation. Residual linear DNA is enzymatically degraded with a novel exonuclease cocktail, leaving highly pure genomic DNA circles. Circularized genomic DNA is treated *in vitro* with ABE RNP complexes to nick the target DNA strand and deaminate adenine bases to inosine (within the base editing window) on the non-target strand at on- and off-target sites (red). Nicked and inosine-containing genomic DNA circles are then processed with endonuclease V which cleaves DNA adjacent to inosines to produce linear DNA with 5’ staggered ends. 5’ overhangs are filled in with Klenow exo-. End-repaired DNA is ligated to sequencing adapters and amplified for high-throughput sequencing. **b,** Dot plot showing the percentage of ABE8e-NRCH:*HBB^S^*-gRNA previously validated off-target sites detected by CHANGE-seq-BE on a subset of enzymatic conditions systematically evaluated during the method development. **c,** Representative alignment of CHANGE-seq-BE reads of the *ABE8e-NRCH:HBB^S^-gRNA* target as visualized by the Integrative Genomics Viewer (IGV). The target adenine base is highlighted in red, and the PAM (NRCH) is highlighted in bold. **d,** Visualization of detected off-targets identified by CHANGE-seq-BE aligned against the intended target site for ABE:gRNA (*ABE8e-NRCH:HBB^S^-gRNA)* complexes targeted against *HBB.* The intended target sequence is shown in the top line, and off-target sites are ordered from top to bottom by CHANGE-seq-BE read count. Mismatches to the intended target sequence are indicated by colored nucleotides, read counts and A-to-I (G) deamination frequencies (%) are shown at the end of each line. For space considerations, this figure shows a list of only the top off-targets detected by CHANGE-seq-BE.

However, a major challenge we encountered was that the addition of the required end-repair step substantially decreased enrichment of editor-modified DNA^21^. To gauge the performance of our initial CHANGE-seq for base editor approach, we quantified the number of validated off-target sites detected for an ABE8e RNP targeting the *HBB* gene (ABE8e-NRCH:*HBB*^S^-gRNA) that we had previously characterized extensively and validated 54 off-target sites. At first, we found that addition of an end repair step to fill in 5’ overhangs and detect base editor modified genomic DNA resulted in high background and detection of only a small fraction (20%) of previously validated off-target sites (**Fig. 1b**). We hypothesized that residual linear DNA molecules could become competent for adapter ligation after end-repair. After systematic optimization of the entire workflow, where we varied circularized genomic DNA exonuclease treatments, base editing reaction conditions, and end repair protocols, we identified an optimized a protocol (**Supplementary Note, Supplementary Protocol**) that consistently detected all 54 previously validated ABE8e-NRCH:*HBB*^S^-gRNA off-target sites, which we called CHANGE-seq-BE (**Fig. 1b)**.

Importantly, CHANGE-seq-BE reads capture a distinct molecular signature of base editing *in vitro,* evidence of A-to-I deamination as inosine bases are converted to guanine after PCR amplification, increasing confidence in sites nominated by our method (**Fig. 1c**). We observed a range of deamination frequencies of 25-48% in top ABE8e-NRCH:*HBB*^S^-gRNA off-targets nominated by CHANGE-seq-BE (**Fig. 1d**).

To evaluate technical reproducibility, we performed independent CHANGE-seq-BE library preparations and found that read counts from technical replicates were strongly correlated (*r* = 0.9814, *r* is Pearson’s correlation coefficient) (**Fig. 2a**), with sites of off-target activity distributed broadly genome-wide (**Fig. 2b**) **(Supplementary Table 1)**. For this gRNA target, the on-target site does not have the highest CHANGE-seq-BE read counts, which is consistent with the lower specificity profile observed.

**Figure 2.**
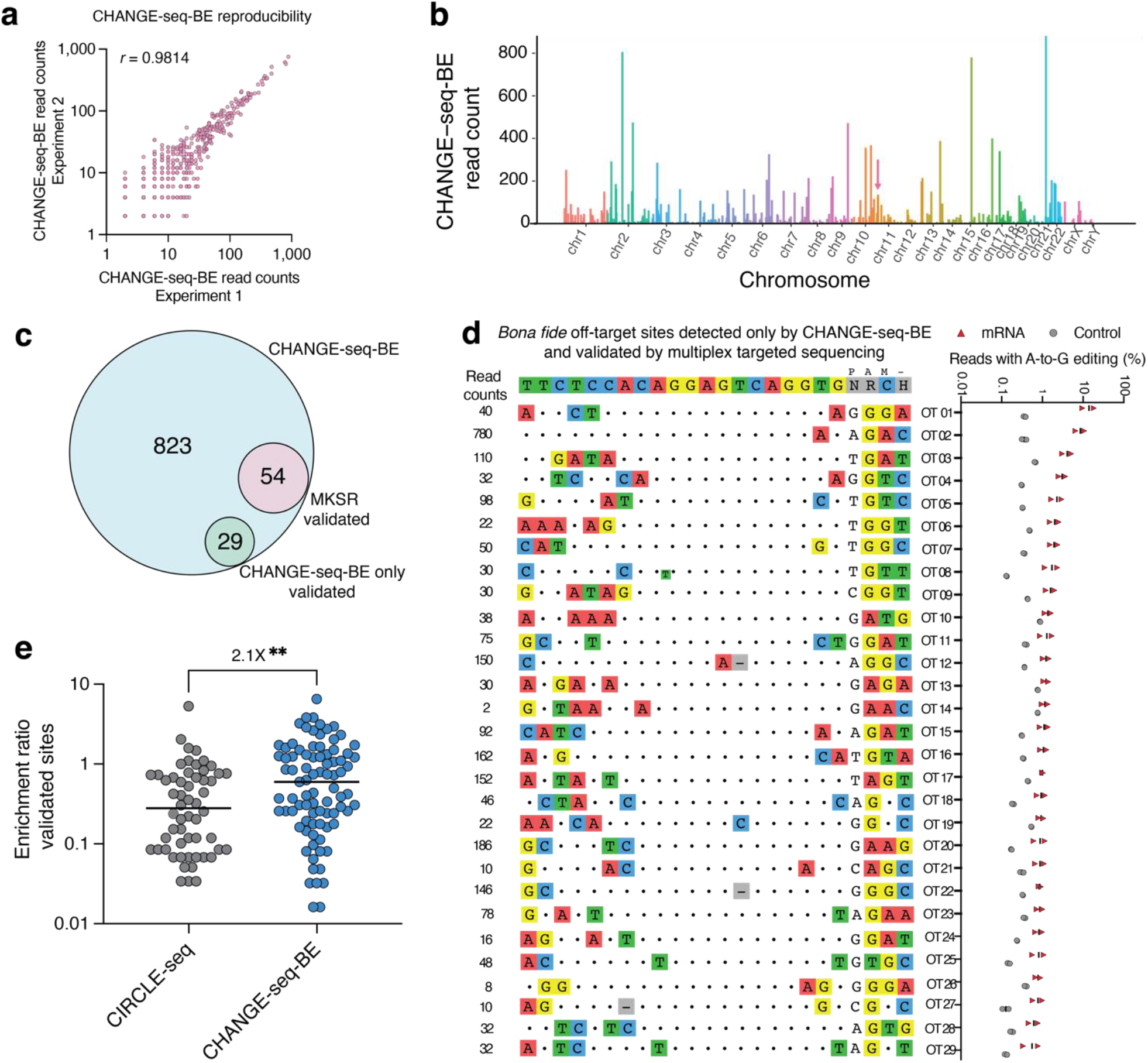
CHANGE-seq-BE sensitively reveals new *bona fide* off-target sites for ABE8e-NRCH:*HBB^S^-gRNA* therapeutic relevant target site. **a,** Scatterplot showing read count correlation for two CHANGE-seq-BE library preparations of ABE8e-NRCH targeting *HBB^S^.* **b,** Manhattan plot showing the genome-wide distribution of CHANGE-seq-BE detected sites for ABE8e-NRCH:*HBB^S^-gRNA*. Red arrow indicates the on-target site. **c,** Venn diagram showing overlap between candidate off-target sites nominated by CHANGE-seq-BE for ABE8e-NRCH:*HBB^S^*-gRNA, and nominated off-target sites for which off-target editing was observed by multiplex targeted sequencing in SCD patient CD34^+^ cells nucleofected with ABE8e-NRCH mRNA. **d,** Dot plot showing the percentage of sequencing reads containing A-to-G base editing within the protospacer positions 4-10 at off-target sites in genomic DNA samples from patient CD34^+^ HSPCs treated with ABE8e-NRCH mRNA (red, n=2), or untreated controls (grey, n=2), P-value < 0.05. **e,** Dot plot showing the enrichment ratio for sites detected by CHANGE-seq-BE using ABE8e, or for CIRCLE-seq using cognate Cas9 nuclease, P-value < 0.05.

To determine whether CHANGE-seq-BE identifies base editor-specific off-target activity not detected by nuclease-specific methods, we compared the sites identified by CHANGE-seq-BE and by CIRCLE-seq using ABE8e-NRCH base editor and Cas9-NRCH nuclease, respectively. We noted that CHANGE-seq-BE and CIRCLE-seq read counts were only weakly correlated (*r* = 0.35) (**Extended Data Fig. 1**), and only 40% of the off-target sites detected by both methods overlap (**Extended Data Fig. 1**), suggesting that ABE8e genome-wide off-target activity quantitatively differs from Cas9 nuclease off-target activity. One possible explanation for some of these differences is the frequency and location of editable adenine bases within on- and off-target sites. Off-targets with mismatches of the editable adenine bases could eliminate base editing activity without affecting nuclease activity.

To confirm that new off-target sites detected exclusively by CHANGE-seq-BE are *bona fide* off-target sites, we performed multiplex targeted sequencing on 131 sites detected exclusively by CHANGE-seq-BE **(Supplementary Table 2)** using the same genomic DNA of CD34^+^ HSPCs treated with ABE8e-NRCH mRNA evaluated in our previous study. We found that CHANGE-seq-BE identified all 54 known off-targets as well as an additional 29 previously unknown *bona fide* off-targets, 53% more than our initial screen with CIRCLE-seq (**Fig. 2c**), with ABE activity ranging from 0.55-13.9% (**Fig. 2d)**. For the *HBB*^S^-gRNA *bona fide* off-target sites identified by CIRCLE-seq using Cas9 nuclease and CHANGE-seq-BE using ABE8e we noted that CHANGE-seq-BE enrichment ratio was significantly higher than CIRCLE-seq, by an average of 2.1-fold (**Fig. 2e**). *HBB*^S^-gRNA *bona fide* off-target activity occurred primarily in intergenic and intronic regions, and six off-targets were located in exons (**Extended Data Fig. 1**). Taken together, our results demonstrate the importance of using sensitive and unbiased base-editor specific off-target discovery methods to comprehensively identify ABE genome-wide off-target activity.

### CHANGE-seq-BE identified unexpected synthetic gRNA contaminants

Safety risks specifically associated with human genome editing products include unintended on- and off-target activity and their long-term consequences. One less obvious but key concern is that contamination in a critical genome editing component such as the gRNA could direct genome editing to unintended on-target sites. Chemically-modified synthetic gRNAs are now commonly used product in genome editing experiments, particularly in primary cells, because of enhanced activity and stability.

Unexpectedly, in initial pilots of CHANGE-seq-BE experiments using research-grade, chemically-synthesized, and HPLC purified gRNAs, we observed CHANGE-seq-BE read counts across multiple targets in base editor treated samples but not in untreated controls, suggesting activity of contaminating synthetic gRNAs (**Fig. 3a**). We detected strong unintended on-target activity for *PDCD1, CD7* and *CIITA* on-target sites in genomic DNA samples treated with *CBLB* gRNA target site; similarly, *PDCD1* and *CD7* reads were found in genomic DNA samples treated with *CIITA* gRNA target site; and *B2M* and *CD7* on-target reads in genomic DNA samples treated with *PDCD1* gRNA target. In hindsight, the most likely source of this cross-contamination during manufacturing was the sequential purification of these gRNAs over the same high-performance liquid chromatography (HPLC) column.

**Figure 3.**
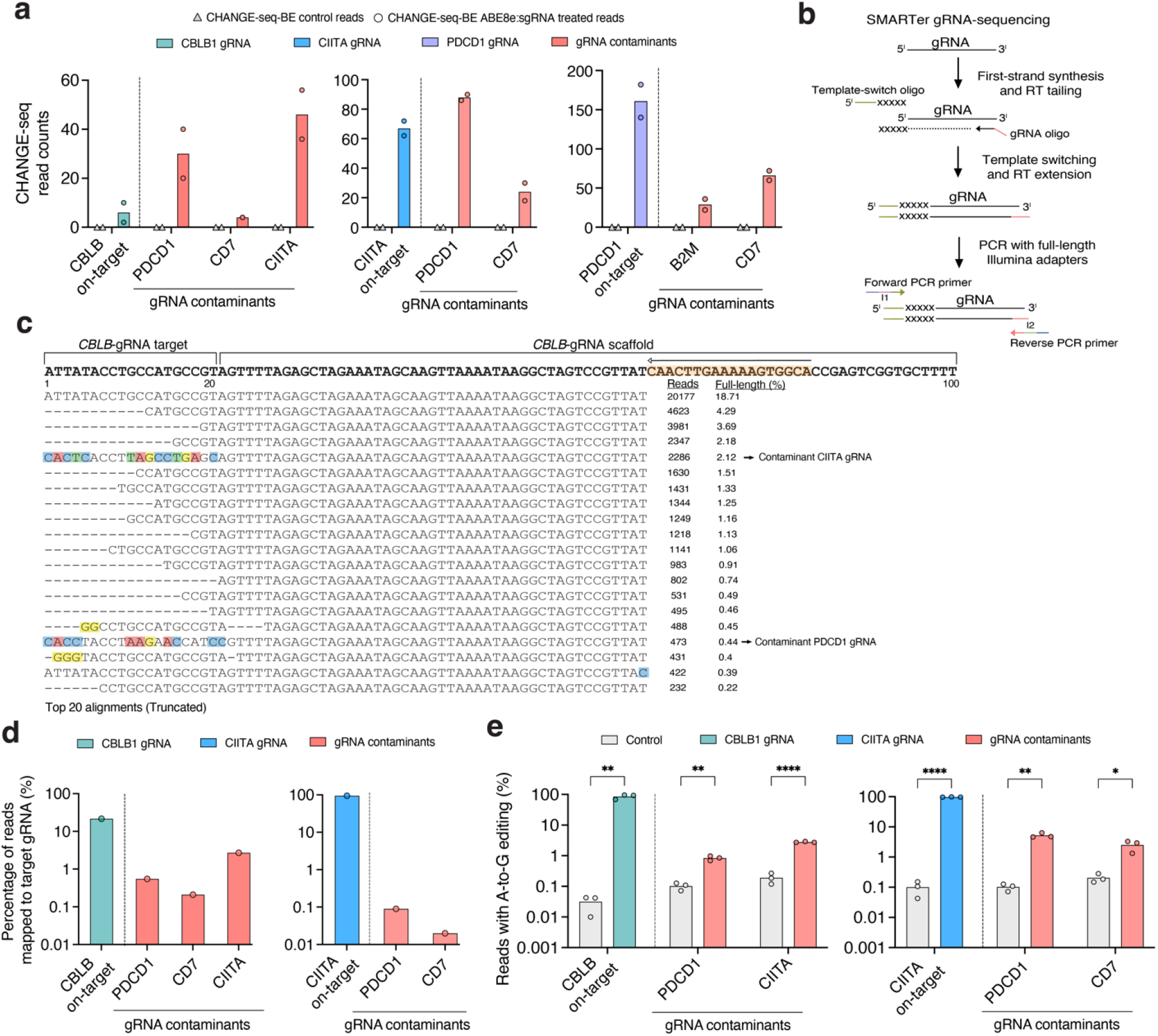
CHANGE-seq-BE identifies activity of synthetic gRNAs contaminants. **a,** Bar plot showing CHANGE-seq-BE read counts for three synthetic chemically modified gRNAs (*CBLB, CIITA* and *PDCD1*) and the unexpected detection of gRNAs contaminants (red). **b,** SMARTer gRNA-sequencing workflow. **c,** Alignment plot for *CBLB*-gRNA target displaying top 20 mapped reads. Mismatched nucleotides in the target sequence are highlighted in colors (blue, green, red, and yellow) representing potential contaminants *CIITA* and *PDCD1*. Arrow highlighted in scaffold region (orange) shows the scaffold specific primer used for first cDNA synthesis. **d,** Bar plot showing the percentage of gRNA RNA-seq reads mapping to target gRNAs, for one specific synthetic gRNAs manufacturers supplier. Activity of gRNA contaminants shown in red. **e,** Bar plot showing the percentage of sequencing reads containing A-to-G mutations within the protospacer positions 4-10 at off-target sites in genomic DNA samples from primary human T cells treated with ABE8e in the mRNA format, and gRNAs targeting *CBLB* or *CIITA*, and the detection of contaminant gRNAs (red) during the manufacture process, P-values for *CBLB* on-target is 0.000657**, gRNA contaminants for *PDCD1* and *CIITA* are 0.000994** and 0.000005****; similarly P-values for *CIITA* on-target was <0.000001****, gRNA contaminants for *PDCD1* and *CD7* are 0.000722** and 0.018647* analyzed compared to controls using multiple unpaired t-tests.

To identify and measure the frequency of contaminating gRNAs, we employed a gRNA sequencing assay based on an established small RNA sequencing technology from Clontech^22^ (**Fig. 3b**). We then evaluated five research-grade gRNA sequences from three synthetic gRNA manufacturer suppliers (A, B and C) for the presence of gRNA contaminants. We detected the presence of contaminating gRNAs in *CBLB* and *CIITA* gRNAs from supplier A (batch 1) (**Fig. 3c-d**), with frequency ranging from 0.09% to 2.71% (**Extended Fig. 2**).

To determine whether the relatively minute amounts of gRNAs contaminants detected *in vitro* by CHANGE-seq-BE activity and gRNA sequencing would be active in cells, we performed targeted sequencing of genomic DNA from primary human T cells edited with ABE8e, with gRNAs manufactured by Supplier A targeting *CBLB* and *CIITA* loci. For cells with intended *CBLB* edits, we additionally evaluated editing at *PDCD1* and *CIITA* on-target sites, and for cells with intended *CIITA* editing, we also evaluated *PDCD1* and *CD7* on-target sites by targeted sequencing. We detected point mutations consistent with adenine base editing in all unintended on-target sites associated with contaminating gRNAs evaluated, with activity ranging from 0.83% to 5.24% (**Fig. 3d**). Our results highlight one feature of using sensitive and unbiased genome-wide methods for base editor off-target discovery is identification of unintended ‘on-target’ activity of contaminant gRNAs. This may be a pervasive issue whenever multiple chemically synthesized gRNAs are purified using the same chromatography supplies and caution is warranted to minimize gRNA synthesis contamination risk, particularly for therapeutic applications.

### CHANGE-seq-BE for therapeutic relevant target sites in human primary cells

To systematically evaluate the genome-wide off-target activity of ABE8e *in vitro*, we performed CHANGE-seq-BE on six therapeutic relevant target sites in human primary T cells (*B2M, CBLB, CD7, CIITA* and *PDCD1*)^23^ or human primary hepatocytes (*PCSK9*)^24^ genomic DNA **(Fig. 4a) (Supplementary Table 1)**. We found that CHANGE-seq-BE read counts were strongly correlated between replicates (*r* > 0.97), highlighting the technical reproducibility of our method (**Extended Data Fig. 3**).

**Figure 4.**
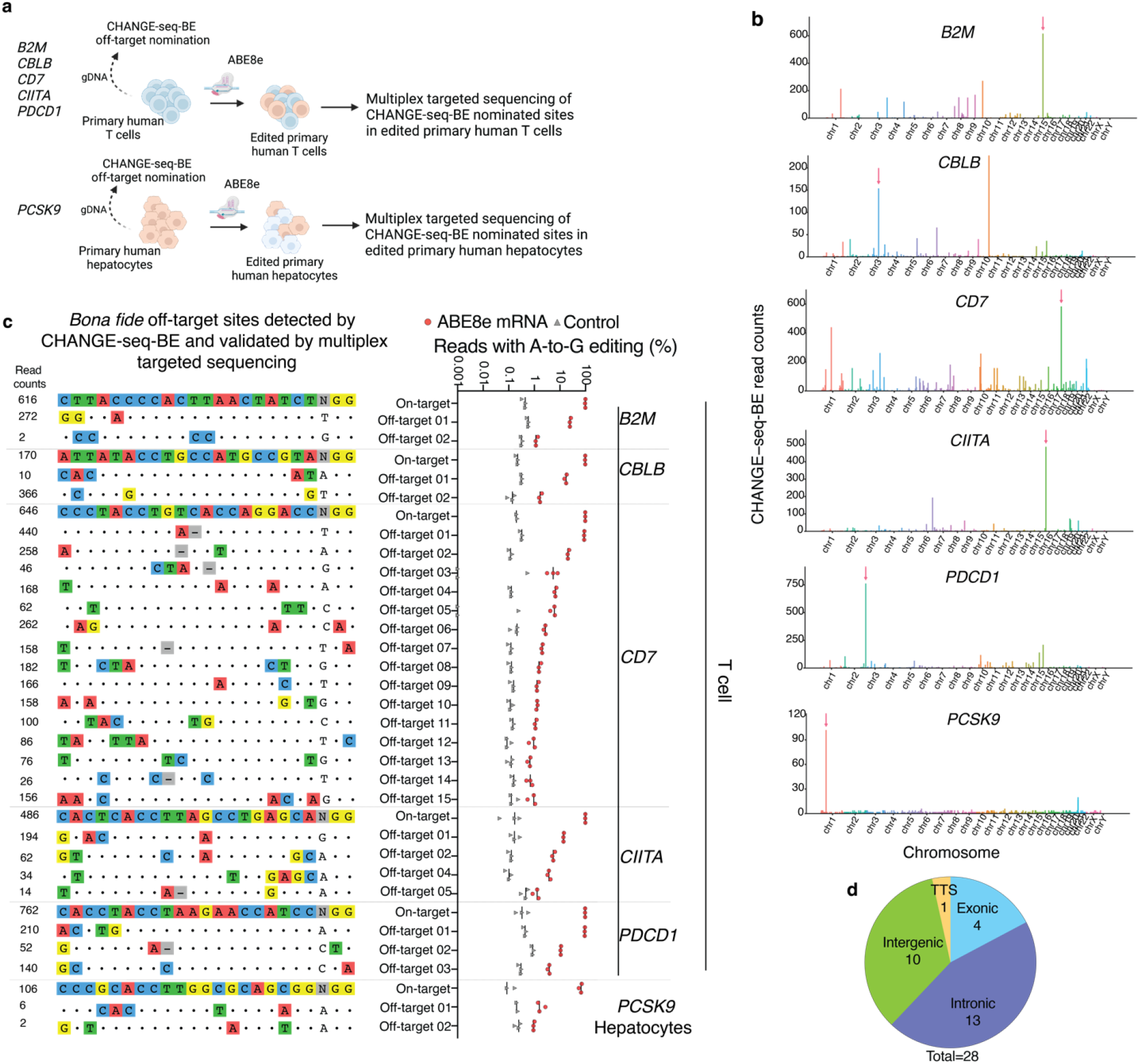
CHANGE-seq-BE detects *bona fide* ABE off-target sites in primary human T cells and hepatocytes. **a**, Schematic illustrating CHANGE-seq-BE off-target nomination in primary human T cells (*B2M, CBLB, CD7, CIITA* and *PDCD1*) and primary human hepatocytes (*PCSK9*). **b,** Manhattan plot showing the genome-wide distribution of CHANGE-seq-BE detected sites. Arrow indicates the on-target site. **c,** Dot plot showing the percentage of sequencing reads containing A-to-G mutations within the protospacer positions 4-10 at off-target sites in genomic DNA samples from primary human T cells (gRNAs targeting *B2M, CBLB, CD7, CIITA* and *PDCD1*) and primary human hepatocytes (gRNA targeting *PCSK9*) treated with ABE8e-NGG mRNA, or untreated controls (n=3). **d,** Predicted genomic features of validated off-target sites (TTS, transcription termination site).

To directly compare ABE8e-NGG and Cas9-NGG off-target activity, we performed CHANGE-seq for four T cell gRNA targets (*B2M, CD7, CIITA* and *PDCD1*) using Cas9 nuclease. For the four therapeutically relevant T cell ABE target sites, CHANGE-seq-BE nominated 6,715 on-and off-target sites, and CHANGE-seq nominated 2,466 sites that were distributed genome-wide (**Fig. 4b**). As expected, although technical replicates for each individual method were very well correlated (r>0.94) (**Extended Data Fig. 3**), CHANGE-seq-BE and CHANGE-seq read counts were less well correlated with each other (*r* = 0.07-0.73) (**Extended Data Fig. 3**), indicating that the biochemical genome-wide off-target profiles of Cas9 and ABE8e are different.

To evaluate the sensitivity of CHANGE-seq-BE to nominate *bona fide* off-target sites, we selected the top off-target sites according to CHANGE-seq-BE read counts (total of 152 off-target sites, varying from 17-27 off-targets per gRNA target site) for analysis by multiplex targeted sequencing in genomic DNA of human primary T cells and human primary hepatocytes treated with ABE8e mRNA. At 152 sites that were successfully amplified and sequenced we confirmed 28 sites (18.4%) **(Supplementary Table 2)** with mutations consistent with ABE8e editing, with activity ranging from 0.92-88.8% (**Fig. 4c).**

To assess the potential functional consequences of off-target activity, we annotated and analyzed the genomic locations of off-targets. Off-target activity occurred predominantly in intergenic and intronic regions, and four were in exons (**Fig. 4d**). In summary, our results demonstrate that CHANGE-seq-BE is a strong predictor of ABE cellular off-target activity in human primary cells and can be used to select candidate target sites for both routine and therapeutic applications.

## Discussion

Here we described CHANGE-seq-BE, which enables the detailed biochemical mapping of base edited genomic DNA and has advantages compared to current methods for defining base editor activity. CHANGE-seq-BE is a way of directly identifying base editor off-target sites that is the first, to our knowledge, to combine desirable qualities of high sensitivity with minimal experimental bias within a single protocol. CHANGE-seq-BE can be used both to rapidly identify highly active and specific base editor and gRNA target combinations.

CHANGE-seq-BE has some key advantages over existing methods. In contrast to biochemical methods that rely on whole genome sequencing of base-editor modified genomic DNA, CHANGE-seq-BE has a substantially reduced rate of observed background reads, enabling it to sensitively identify ABE off-target sites and be ∼20-fold more sequencing efficient than existing alternative genome-wide unbiased methods. The method requires ∼25 million sequencing reads, therefore it is accessible to most labs and is amenable to automation and high-throughput scaling. Importantly, it directly measures base editor off-target activity and is therefore highly relevant for IND-enabling characterization of clinical genome editing approaches.

One limitation of our approach is that CHANGE-seq-BE libraries currently require higher sequencing depth to achieve similar on-target read counts as CHANGE-seq libraries for nucleases. This may be associated with guide-independent deamination or other technical factors, which we plan to explore and optimize further.

We anticipate that CHANGE-seq-BE should also be readily adaptable to cytosine base editors, such as recently engineered cytosine base editor variants of TadA^25–27^. We have previously automated CHANGE-seq on a laboratory automation platform and expect that CHANGE-seq-BE should be similarly automatable.

An ongoing challenge remains functional characterization of potential base editor off-target sites. For example, it is not technically straightforward to perturb off-target sites individually without affecting other closely related sites, and thus difficult to de-risk them without other confounding base edits. Thus, although CHANGE-seq-BE represents an important advance to nominating base editor off-target sites, complementary approaches will be required to dissect their function.

Overall, we anticipate that CHANGE-seq-BE will become an essential way to characterize the genome-wide off-target activity of base editors for both life sciences research and clinical applications.

## Methods

### Isolation of human primary T cells

Research-consented and deidentified peripheral blood mononuclear cells (PBMCs) were obtained commercially (Key Biologics); CD4^+^/CD8^+^ T cells were purified using magnetic separation on a CliniMACS Plus instrument. Cells were washed in CliniMACS buffer with 0.5% HSA and resuspended in 190 ml of IVIG (Intravenous Immunoglobulin), followed by incubation for 15 minutes. Subsequently, cells were incubated and labeled with CD4 and CD8 microbeads (Miltenyi Biotec). Next, two washes were performed using CliniMACS buffer with 0.5% HSA, followed by CD4^+^/CD8^+^ cells selection. The percentage of CD3^+^, CD4^+^, CD8^+^, and CD19^+^ cells in the selected population was determined by flow cytometry as quality control metrics.

### ABE8e (NGG and NRCH) expression and purification

ABE8e-NGG and ABE8-NRCH were expressed and purified as previously described^12,28^, at the Protein Production Core Facility at St. Jude. Briefly, the expression plasmids (pRha-ABE8e-NRCH, Addgene #165417; and pABE8e-protein, Addgene #161788) were transformed into BL21Start DE3 competent cells (Thermo), and colonies were picked for growth in terrific broth (TB) with kanamycin (25 mg/ml) at 37 °C. After overnight growth, 1l of pre-warmed TB was inoculated with the culture at a starting OD600 of ∼0.05. Cells were kept at 37 °C and shaking at 250 rpm until the OD600 reached 1.5. Next, the cells were cold-shocked in an ice-water slurry, following which L-rhamnose (0.8%) was added to induce expression. Cells were then kept shaking at 18 °C for 24 h for protein expression. Following induction, cells were pelleted and fand resuspended in 30 ml cold lysis buffer (1 M NaCl, 100 mM Tris-HCl pH 7.0, 5 mM TCEP, 10% glycerol, with 5 tablets of complete, EDTA-free protease inhibitor cocktail (Millipore Sigma). Cells were passed three times through a homogenizer (Avestin Emulsiflex-C3) at ∼18,000 psi to lyse. Cell debris was pelleted for 20 min using 20,000g centrifugation at 4 °C. Supernatant was collected and spiked with 40 mM imidazole, followed by a 1 h incubation at 4 °C with 1 ml Ni-NTA resin slurry (G Bioscience). Protein-bound resin was washed twice with 12 ml lysis buffer in a gravity column at 4 °C. Protein was eluted in 3 ml elution buffer (300 mM imidazole, 500 mM NaCl, 100 mM Tris-HCl pH 7.0, 5 mM TCEP, 10% glycerol). Eluted protein was diluted in 40 ml low-salt buffer (100 mM Tris-HCl, pH 7.0, 5 mM TCEP, 10% glycerol) just before loading into a 50-ml Akta Superloop for ion exchange purification on an Akta Pure25 FPLC. Ion exchange chromatography was conducted on a 5-ml GE Healthcare HiTrap SP HP pre-packed column. After washing the column with low-salt buffer, we flowed the diluted protein through the column to bind. The column was then washed in 15 ml low-salt buffer before being subjected to an increasing gradient to a maximum of 80% high-salt buffer (1 M NaCl, 100 mM Tris-HCl, pH 7.0, 5 mM TCEP, 10% glycerol) over the course of 50 ml, at a flow rate of 5 ml per minute. One-ml fractions were collected during this ramp to high-salt buffer. Peaks were assessed using SDS–PAGE to identify fractions that contained the desired protein, which were concentrated first using an Amicon Ultra 15-ml centrifugal filter, followed by a 0.5-ml 100-kDa cutoff Pierce concentrator. Concentrated protein was quantified using a BCA assay (Thermo Fisher).

### Cell culture

Human primary T cells were cultured in X-Vivo 15 media (Lonza) supplemented with 10% human heat-inactivated serum (HSA) (Fisher), 10 ng/ml IL-7 (Miltenyi), and 10 ng/ml IL-15 (Miltenyi) at 37 °C with 5% CO_2_. T cells were stimulated with MACS GMP T-cell TransAct polymeric nanomatrix (Miltenyi) for 3 days according to the manufacturer’s instructions prior to transfection. Human primary hepatocytes were cultured in Upcyte Hepatocyte High Performance Medium (BioIVT) with supplement A (BioIVT) and L-glutamine (BioIVT), in plates coated with collagen type I (Sigma) at 37 °C with 5% CO_2_.

### CHANGE-seq-BE

CHANGE-seq-BE was performed on genomic DNA isolated from human CD4^+^/CD8^+^ T cells or human primary hepatocytes (BioIVT). Genomic DNA was isolated using Gentra PureGene Tissue Kit (Qiagen) and quantified by Qubit fluorimetry (Invitrogen). Purified genomic DNA was tagmented with a custom Tn5-transposome to an average length of ∼500 bp, gap repaired with HiFi HotStart Uracil+ Ready Mix (Kapa) and Taq DNA ligase (NEB), and treated with a mixture of USER enzyme and T4 polynucleotide kinase (NEB). DNA was circularized at a concentration of 5 ng/µl with T4 DNA ligase (NEB), and treated with a cocktail of exonucleases, Lambda exonuclease (NEB), Exonuclease I (NEB), Plasmid-Safe ATP-dependent DNase (Lucigen) and Exonuclease III (NEB), to enzymatically degrade remaining linear DNA molecules, followed by dephosphorylation with Quick SAP (NEB). gRNAs were re-folded prior ABE8e:gRNA complexation, and a ABE8e:gRNA ratio of 1:3 was used to ensure full ribonucleoprotein complexation. *In vitro* deamination reactions were performed in a 50 µl volume with deamination buffer (50 mM TRIS-HCl (pH 8), 25 mM KCl, 2.5 mM MgSO_4_, 0.1 mM EDTA, 10% glycerol, 2 mM DTT, 10 µM ZnCl_2_, 300 nM of ABE, 900 nM of gRNA (listed in **Supplementary Table 3**), and 125 ng of circularized DNA. ABE8e deaminated DNA products were treated with proteinase K (NEB), followed by Endonuclease V (NEB) treatment. Endonuclease V-treated DNA molecules were end-repair with Klenow exo-(NEB), A-tailed, ligated with a hairpin adapter (NEB), treated with USER enzyme and amplified by PCR using HiFi HotStart Uracil+ Ready Mix (Kapa/Roche). Completed libraries were quantified by qPCR using Kapa Library Quantification kit (Kapa Biosystems) and sequenced with 151-bp paired-end reads on an Illumina NextSeq 550 instrument. A detailed user protocol for CHANGE-seq-BE is provided (**Supplementary Protocol**).

### CHANGE-seq

CHANGE-seq was performed as previously described. Briefly, genomic DNA from human primary T cells was isolated using Gentra Puregene Kit (Qiagen) according to manufacturer’s instructions. CHANGE-seq was performed as previously described^21^. Briefly, purified genomic DNA was tagmented with a custom Tn5-transposome to an average length of 400 bp, followed by gap repair with Kapa HiFi HotStart Uracil+ DNA Polymerase (KAPA Biosystems) and Taq DNA ligase (NEB). Gap-repaired tagmented DNA was treated with USER enzyme (NEB) and T4 polynucleotide kinase (NEB). Intramolecular circularization of the DNA was performed with T4 DNA ligase (NEB) and residual linear DNA was degraded by a cocktail of exonucleases containing Plasmid-Safe ATP-dependent DNase (Lucigen), Lambda exonuclease (NEB) and Exonuclease I (NEB). *In vitro* cleavage reactions were performed with 125 ng of exonuclease-treated circularized DNA, 90 nM of SpCas9 protein (NEB), NEB buffer 3.1 (NEB) and 270 nM of gRNA, in a 50 μL volume. Cleaved products were A-tailed, ligated with a hairpin adaptor (NEB), treated with USER enzyme (NEB) and amplified by PCR with barcoded universal primers NEBNext Multiplex Oligos for Illumina (NEB), using Kapa HiFi Polymerase (KAPA Biosystems). Libraries were quantified by qPCR (KAPA Biosystems) and sequenced with 151 bp paired-end reads on an Illumina NextSeq instrument. CHANGE-seq data analyses were performed using open-source CHANGE-seq analysis software (https://github.com/tsailabSJ/changeseq/dev).

### CHANGE-seq-BE analysis

Adapter trimming was performed using cutadapt (v1.18) with parameters of “--overlap 40 -e 0.15” to remove Tn5 adapter (CTGTCTCTTATACACATCTACGTAGATGTGTATAAGAGACAG). Then paired-end reads were mapped using bwa mem with default parameters. CHANGE-seq-BE paired-end reads should be in an outward orientation with some degree of self-overlapping bases (default min_overlap=0 and max_overlap=15). Reads were filtered based on the above criteria. The start positions of the valid reads were then tabulated, and genomic intervals enriched in nuclease-treated samples were identified. The interval and 30 bp of flanking reference sequence on either side were searched for potential nuclease-induced off-target sites with an edit distance of less than or equal to six, allowing for gaps. Deamination events can be A-to-G conversion if the off-target’s strand is forward or C-to-T conversion if the off-target’s strand is reverse. Deamination-containing reads were counted if the deamination events occurred close to +2 of the read start position (allowing for +/- 2 bp flanking). The percentage of deamination was calculated as the ratio between number of deamination-containing reads and the total number of valid reads assigned to the off-target.

### Cell transfection

Transfections of human primary T cells were performed with 2 µg of ABE8e mRNA and 1 µg of synthetic chemically modified gRNA. mRNA:gRNA were added directly to 1×10^6^ cells resuspended in 20 µl of P3 solution and nucleofected with pre-programmed pulse EO-115 in 4D-Nucleofector^TM^ System (Lonza). After nucleofection, cells were recovered in X-Vivo 15 media with 20% human heat-inactivated serum (Fisher), 10 ng/ml IL-7 (Miltenyi), and 10 ng/ml IL-15 (Miltenyi). After 5 days cells were harvested for genomic DNA purification using Agencourt DNAdvance (Beckman Coulter). Transfection of human primary hepatocytes were performed using Lipofectamine MessengerMAX Transfection Reagent (Thermo Fisher). Briefly, 1.5 x10^5^ hepatocytes were seeded in a 6-well plate coated with collagen type I. After 24 h, cells were transfected with 2.5 µg of ABE8e mRNA, 2.5 µg of synthetic chemically modified gRNA (*PCSK9*) and 7.5 µl of Lipofectamine Messenger MAX, following the manufacturers recommendations. After three days cells were harvested for genomic DNA purification using Agencourt DNAdvance (Beckman Coulter).

### Multiplex targeted amplicon sequencing

To determine the A-to-G conversion frequency at CHANGE-seq-BE identified sites, on- and off-target sites were amplified from genomic DNA from edited cells or unedited controls using rhAmpSeq system (IDT), in triplicates, with primers listed in **Supplementary Table 2**. Note that for the validation of the newly CHANGE-seq-BE detected sites for the MKSR targeted, we used the same genomic DNA of CD34^+^ treated with ABE8e-NRCH:*HBB*^S^-gRNA mRNA described in our previous study^12^. Sequencing libraries were generated according to manufacturer’s instructions. Completed libraries were quantified by qPCR using Kapa Library Quantification kit (Kapa Biosystems) and sequenced with 151-bp paired-end reads on an Illumina MiSeq or MiniSeq instrument.

### Targeted sequencing

To determine the A-to-G conversion frequency at intended and unintended on-target sites, primary human T cells were nucleofected with ABE8e mRNA and *CBLB* or *CIITA* gRNAs. On-target sites (*CBLB*, *CIITA*, *B2M*, *CD7* or *PDCD1*) were amplified from T cell genomic DNA using 2X Phusion Hot Start Flex Master Mix (NEB) (primers described in **Supplementary Table 3**) and 100 ng of genomic DNA as the input for each PCR. PCR products were purified with SPRI magnetic beads, normalized, and 10 ng of each amplicon were used for the second PCR to add complete Illumina adapters and indexes. Completed libraries were quantified by qPCR using Kapa Library Quantification kit (Kapa Biosystems) and sequenced with 151-bp paired-end reads on an Illumina MiSeq or MiniSeq instrument.

### Quantification of base-editing mutations at evaluated on-target sites and guide dependent off-target sites

Adenine base editing frequency for each position in the protospacer was quantified using CRISPRessoPooled (v2.0.41) with ‘quantification_window_size_10 - quantification_window_center-10-base_editor_output-conversion_nun_fromA - conversion_nuc_toG’. The genomic features of all bona fide off-target sites detected by CHANGE-seq-BE were annotated using HOMER (v.4.10). (Ref). To calculate the statistical significance of off-target editing for the ABE8e mRNA treatments compared to control samples, we applied a chi-square test (two replicates). The contingency table was construct using the number of edited reads and the number of unedited reads in treated and control groups. The FDR was calculated using the Benjamini-Hochberg method. The reported significant off-targets were called on the basis of FDR < 0.05 and difference in editing frequency between treated and untreated control >0.5% for at least one treatment. The custom code used to conduct off-target quantification and the statistical analysis is available to download at https://github.com/tsailabSJ/MKSR_off_targets.

### gRNA sequencing

To generate gRNA-seq libraries, we repurposed the SMARTer smRNA-seq kit (Takara). First, cDNA synthesis was performed with 1-2 ng of gRNA template using 1 μl of custom-tailed scaffold specific primer (10μM) 5’ GTGACTGGAGTTCAGACGTGTGCTCTTCCGATCT <u>TGCCACTTTTTCAAGTTG</u> 3’. Incubate the reaction at 72°C for 3 minutes and 4°C for 2 minutes to anneal custom-tailed scaffold primer at the 3’ end of the scaffold gRNA. Reverse transcription was performed according to the manufacturer’s instructions. After cDNA synthesis, a library enrichment PCR was performed with barcoded universal primers with following PCR conditions, denaturation - 98°C for 1:00; annealing (17x cycles) - 98°C 60°C 68°C for 0:10 0:05, 0:10; hold 10°C. Final libraries were purified with Nucleospin PCR Clean up kit (Takara) and quantified by Qubit HS (Thermo Scientific). The pooled library with indexes was loaded at 2.2nM (+10% phix) and sequenced with 101bp paired-end reads on an Illumina MiSeq instrument.

### gRNA sequencing analysis

Paired-end reads were merged using FLASH with an expected fragment length of 100bp. Scaffold specific primer (3’-end) was trimmed using cutadapt (v4.1) and read with length than 20bp were removed. The percentage of target gRNA and full length product (target gRNA + scaffold sequence) were calculated based on an exact string match. Merged reads were aligned to the expected full length product using bowtie2 to identify indels in the gRNA sequencing data. Source code is available at: https://github.com/YichaoOU/gRNA_sequencing.

---

**Extended Figure 1.**
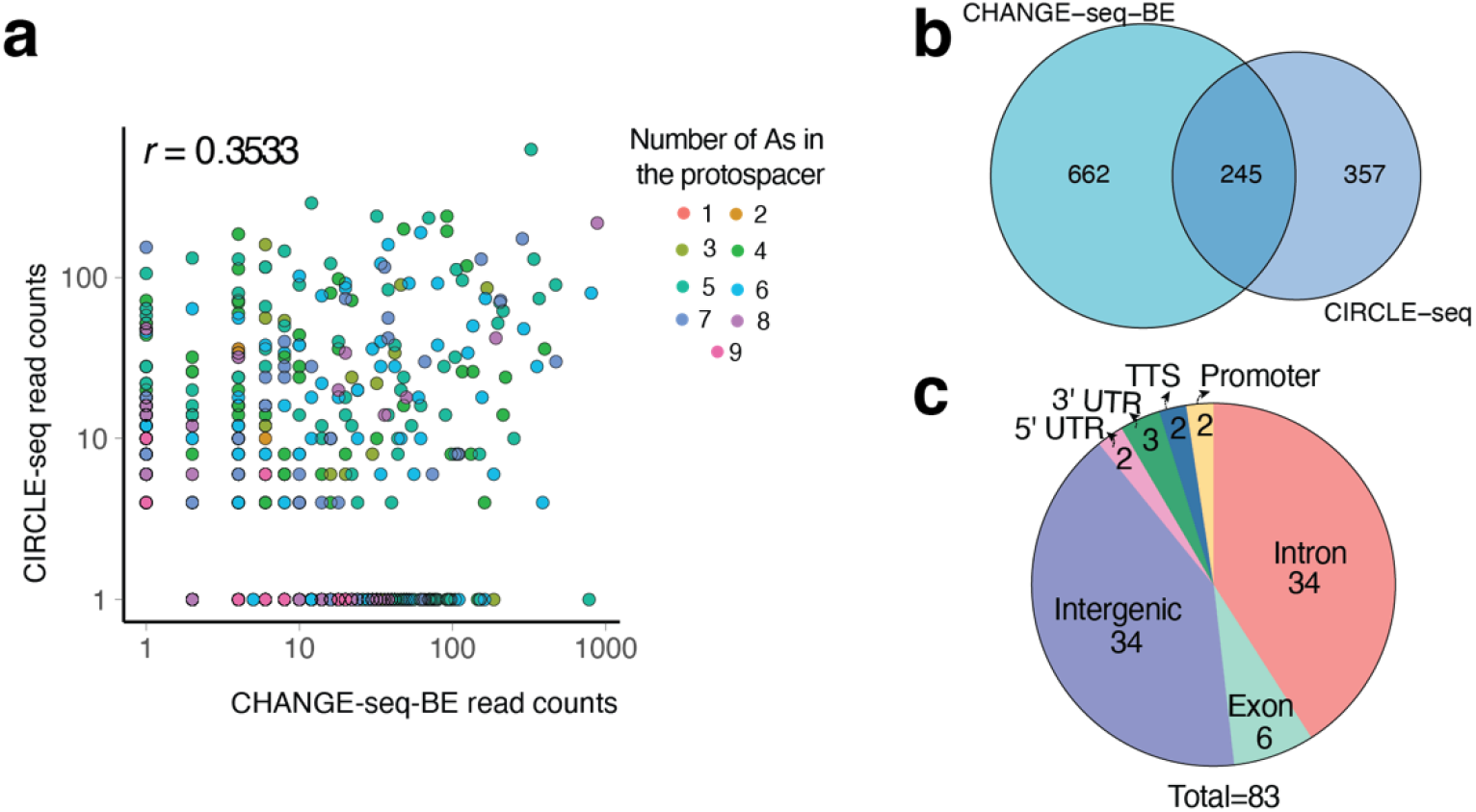
CHANGE-seq-BE detects new off-target sites for ABE8e-NRCH:*HBB*^S^-gRNA. **a,** Scatterplot showing correlation of CHANGE-seq-BE read counts using ABE8e-NRCH and CIRCLE-seq using cognate Cas9-NRCH, colored by the number of adenines in the protospacer positions. **b,** Venn diagram depicting the overlap of off-target sites detected by CHANGE-seq-BE ABE8e-NRCH and CIRCLE-seq Cas9-NRCH for the *HBB^S^*-gRNA target. **c,** Pie chart of genomic annotations of all validated ABE off-target sites for the ABE8e-NRCH:*HBB*^S^-gRNA target site.

**Extended Figure 2.**
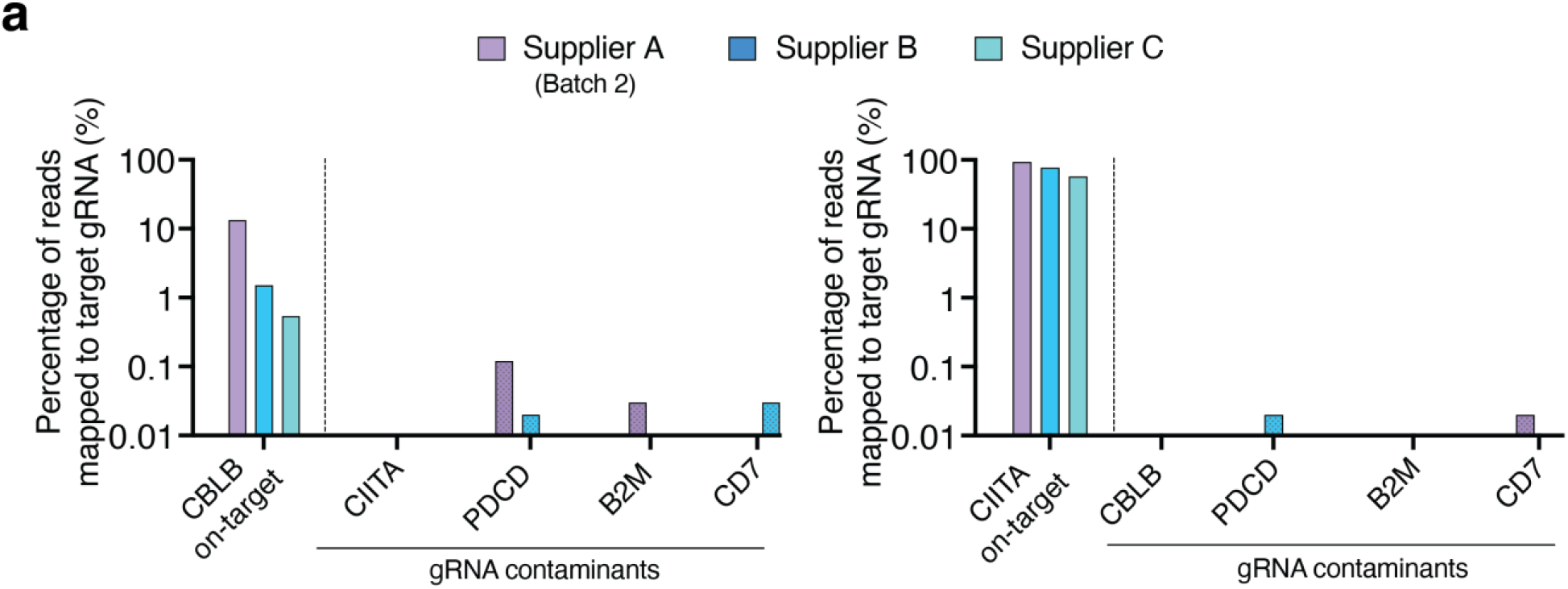
gRNA sequencing detects potential gRNA contaminants in target gRNAs. Bar plot showing the percentage of gRNA sequencing reads mapping to target gRNAs for the three synthetic gRNAs manufacturers suppliers.

**Extended Figure 3.**
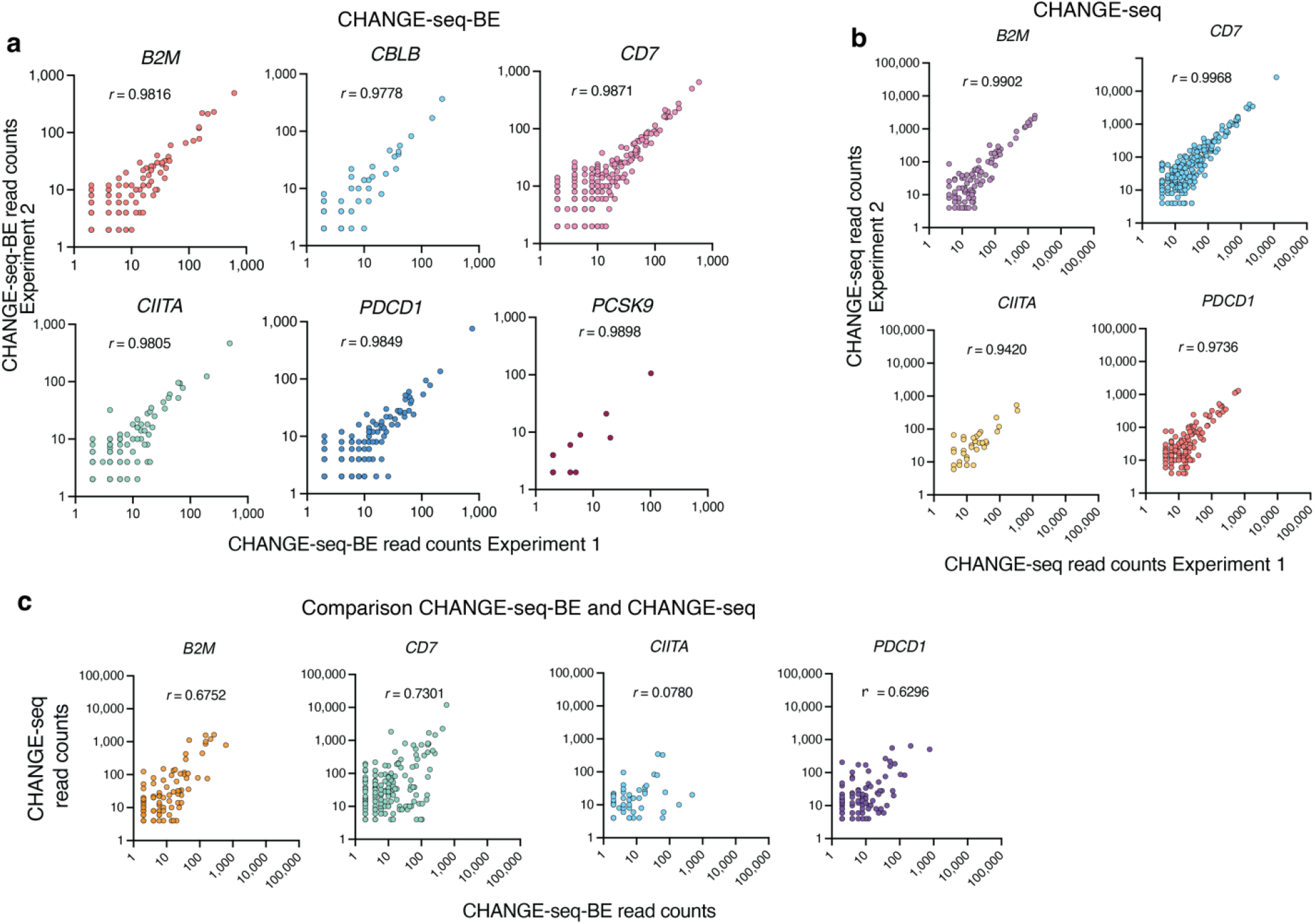
*In vitro* off-target profile of ABE and Cas9. **a,** Scatterplot showing read counts correlation for two independent CHANGE-seq-BE (ABE8e) technical replicates. **b,** Scatterplot showing read counts correlation for two independent CHANGE-seq (Cas9) technical replicates. **c,** Scatterplot showing read counts correlation for ABE8e and Cas9 for five T cell relevant target sites.

## Acknowledgments

We thank D. Liu and G. Newby for ABE8e-NGG and ABE8e-NRCH protein expression plasmids, W. Wang from St. Jude Protein Production Core Facility for recombinant Tn5 and recombinant ABE8e expression and purification, J. Yen and M. Weiss for ABE8e-NRCH-HBB edited CD34 cells genomic DNA. This work was supported by St. Jude Children’s Research Hospital and ALSAC, National Institutes of Allergy and Infectious Diseases award U01AI176471 (to SQT), National Heart Lung and Blood Institute award U01HL163983 (to SQT), St Jude Children’s Research Hospital Collaborative Research Consortium on Novel Gene Therapies for Sickle Cell Disease and the Doris Duke Charitable Foundation (2020154).

## Author contributions

C.R.L and S.Q.T conceived of and designed the study. C.R.L, V.K, E.U and G.L performed experiments. Y.L performed computational analysis. C.R.L and S.Q.T wrote the paper with input from all authors.

