## Supplementary Information for "CHANGE-seq-BE enables simultaneously sensitive and unbiased *in vitro* profiling of base editor genome-wide activity"

### Supplementary note

#### CHANGE-seq-BE optimization

Previously, we developed CHANGE-seq (Circularization for high-throughput analysis of nuclease genome-wide effects by sequencing)[1], a sensitive, unbiased, and high-throughput method for defining CRISPR-Cas9 off-target activity in vitro that has been widely adopted by academic and industries laboratories. To leverage CHANGE-seq to define the genome-wide off-target activity of other genome editors, such as adenine base editors, additional enzymatic steps are required. *In vitro*, adenine base editors RNP complexes nick the target DNA strand and deaminate adenine bases to inosine (within the base editing window) on the non-target strand at on- and off-target sites. Nicked and inosine-containing genomic DNA are further processed with endonuclease V, which cleaves DNA adjacent to inosines to produce linear DNA with 5' staggered ends. However, in the current CHANGE-seq workflow, the newly DNA ends generated by Cas9 cleavage of circularized DNA are directly A-tailed, and sequencing adapters are ligated, bypassing the requirement of an end repair step, as the majority of the DSBs generated by Cas9 in the DNA produces blunt ends. Previously, we found that the simple addition of an end repair step into CHANGE-seq workflow substantially reduced the sensitivity of the method, suggesting the persistence of some linear DNA, presumably in a very low concentration.

To stringently eliminate remaining linear DNA molecules after genomic DNA circularization, we evaluated a set of exonucleases, including Plasmid-safe (Lucigen), E.coli exonuclease I (NEB), Lambda exonuclease (NEB), Exonuclease III (NEB), T5 exonuclease (NEB), T7 exonuclease (NEB), Bal31 exonuclease (NEB) and their combinations thereof, as well as enzymatic reaction buffers. We found that the combination of Plasmid-Safe, Lambda exonuclease, E. coli Exonuclease I and Exonuclease III in Ampligase buffer (Lucigen) improved CHANGE-seq on-target enrichment when end repair was introduced into the workflow. Next, we then tested two end repair strategies using Kapa end repair mix (fill in of 5' overhangs and removal of 3' overhangs to form blunt ends) or Klenow fragment (3'→5' exo negative) (NEB) (fill in of 5' overhangs only, to form blunt ends). We found that filling in 5' overhangs only with Klenow fragment (3'→5' exo negative) outperformed the Kapa end repair mix treatment. We also evaluate the addition of a Quick CIP (calf intestinal alkaline phosphatase) treatment after stringent exonuclease selection of circularized DNA to prevent adapter ligation in existing blunt DNA ends. We found that the association of Quick CIP treatment of circularized DNA before *in vitro* deamination, and Klenow exo- treatment to fill in 5' ends generated by ABE and Endonuclease V activity further reduced sequencing background and enriched CHANGE-seq read counts.

After defining the ideal combination of exonucleases to compose a stringent cocktail of enzymes for circular DNA selection and the best end repair strategy, we then sought to further improve the sensitivity of the method exclusively for the enrichment of ABE modified genomic DNA molecules in the CHANGE-seq workflow optimized for base editors (CHANGE-seq-BE). We evaluated several conditions for deamination, such as deamination buffer, deamination time and RNP concentration. Additionally, we tested different concentrations of Endonuclease V and two DNA polymerases for PCR enrichment of DNA containing inosine. To test all these conditions, we leveraged the ABE8e-NRCH:*HBB*$^{S}$-sgRNA, a target site and editor that we extensively characterized previously[2], with 54 known bona fide cellular ABE8e off-target sites. The final selected conditions were determined based on CHANGE-seq-BE enrichment (read counts) and the percentage of known ABE8e-NRCH:*HBB*$^{S}$-sgRNA bona fide off-targets (54 sites) retrieved by the new CHANGE-seq-BE workflow.

#### Supplementary Protocol: CHANGE-seq-BE

**REAGENTS**

- Gentra Puregene Tissue Kit (Qiagen, cat.no. 158667)
- IDTE pH 8.0 (1X TE Solution) (Integrated DNA Technologies, cat.no. 11050204)
- Tn5 transposase (St. Jude Protein Core. Recombinant Tn5 is expressed and purified according to Picelli *et al* (2014)[7], using Rosetta$^{TM}$ 2(DE3) pLysS Competent Cells)
- Proteinase K (NEB, cat.no. P8107S)
- KAPA HiFi HotStart Uracil+ ReadyMix (250 x 50 µl reactions) (Kapa Biosystems, cat.no. KK2802)
- Taq DNA ligase (NEB, cat.no. M0208L)
- T4 Polynucleotide Kinase (PNK) (New England BioLabs, cat.no. M0201L)
- T4 DNA Ligase (New England BioLabs, cat.no. M0202L)
- 10X T4 DNA ligase Buffer (New England BioLabs), supplied with T4 DNA Ligase
- USER Enzyme (New England BioLabs, cat.no. M5505L)
- Exonuclease I (*E. coli*) (New England BioLabs, cat.no. M0293L)
- Lambda Exonuclease (New England BioLabs, cat.no. M0262L)
- Exonuclease III (E. coli) (New England BioLabs, cat.no. M0206L)
- Plasmid-Safe ATP-dependent DNase (Epicentre, cat.no. E3110K)
- 10X Ampligase Buffer (Lucigen, cat.no. A1905B)
- Plasmid-Safe 10X Reaction Buffer (Epicentre), supplied with Plasmid-Safe ATP-dependent DNase
- 25mM ATP solution (Epicentre), supplied with Plasmid-Safe ATP-dependent DNase
- Quick CIP (NEB, cat.no. M0525L)
- ABE8e (St. Jude Protein Core. ABE8e is expressed and purified according to Huang, Newby and Liu (Nat. Protocols, 2021))
- Endonuclease V (NEB, cat.no. M0305S)
- Klenow Fragment (3' -> 5' exo-) (NEB, cat.no. M0212L)
- dNTPs (NEB, cat.no. N0447L)
- NEBNext dA-tailing Module (NEB, cat.no. E6053)
- NEBNext Ultra II Ligation Module (NEB, cat.no. E7595)
- PEG/NaCl SPRI solution (Kapa Biosystems, cat. no. 07961928001)
- NEBNext® Multiplex Oligos for Illumina® (Dual Index Primers Set 1) (New England BioLabs, cat.no. E7600S)
- NEBNext adapter for Illumina (New England BioLabs), supplied with NEBNext® Multiplex Oligos for Illumina®
- Qubit dsDNA BR Assay Kit (Thermo Fisher Scientific, cat.no. Q32853)
- Qubit dsDNA HS Assay Kit (Thermo Fisher Scientific, cat.no. Q32854)
- Qubit assay tubes (Thermo Fisher Scientific, cat.no. Q32856)
- NextSeq kit 300-cycles (mid or high output kit) (Illumina)
- Flow Cell, supplied with NextSeq® Reagent Kit
- NextSeq Buffer, supplied NextSeq® Reagent Kit
- PhiX Control V3 KIT (Illumina, cat.no. FC-110-3001)
- Sodium hydroxide solution, volumetric, 1 M NaOH (1N) (Sigma-Aldrich, cat.no. 71463-1L
- North Alcohol Wipes (Dynarex, cat.no. 19-014-855)
- VWR Lens Cleaning Tissue (VWR, cat.no. 52846-001)
- EDTA 0.5 M (Thermo Fisher Scientific, cat.no. 15575020)
- Ethanol (Sigma, cat.no. E7023)
- Tween-20 (Sigma-Aldrich, cat.no. P7949)
- Sera-Mag Magnetic Beads; Carboxyl, Speedbeads; hydrophobic; 5 solids (Fisher/GE, cat.no. 9981123)
- Guanidine thiocyanate (Sigma, cat.no. G9277)
- Sodium Chloride 5 M Sterile (Fisher, cat.no. 50146927)
- TRIS Buffer 1.0 M solution, pH 8.0 (Fisher, cat.no. 50146868)
- Polyethylene Glycol 8000 (Fisher, cat.no. 507516674)
- Lib Quant Kit (Illumina/Uni) (Kapa Biosystems, cat.no. KK4824)
- N,N-Dimethylformamide (Sigma, D4551-250ML)

- TAPS (Sigma, T5130)
- Magnesium chloride hexahydrate (Sigma, M9272-500G)

**REAGENT SETUP**

**Resuspend the CHANGE-seq/CHANGE-seq-BE custom transposon oligonucleotides (oCRL225 and oCRL226)** Resuspend the oligonucleotides to 100 µM in TE pH 8.0. Keep the resuspended oligonucleotides at -20°C.

**oCRL225** /5Phos/ACG/ideoxyU/AGATGTGTATAAGAGACAG

**oCRL226** /5Phos/CTGTCTCTTATACACATCTACGT

**Anneal CHANGE-seq custom transposon (oCRL225 and oCRL226)** Mix the oligonucleotides as follows:

| Component | Volume (µl) |
|---|---|
| oCRL225 100 µM | 50 |
| oCRL226 100 µM | 50 |
| Total | 100 |

On a thermocycler, set up the follow annealing program: 95°C for 5 min, -1°C/min for 70 cycles, hold at 4°C.
After annealing, add 100 µl of TE pH 8.0 to bring the concentration of the annealed oligonucleotides to 25 µM. Keep the annealed oligonucleotides at -20°C. The annealed adapters will be used for transposome assembly.

**2X Tn5 dialysis buffer** 100 mM Hepes-KOH, pH 7.2, 0.2 M NaCl, 0.2 mM EDTA, 2 mM DTT, 0.2% Triton X-100, 20% glycerol.

**Transposome assembly** Perform the transposome assembly as follows:

| Component | Volume (µl) |
|---|---|
| Tn5 (1.85 mg/ml) | 360 |
| Annealed oCRL225/oCRL226 (25 µM) | 150 |
| 2X Tn5 dialysis buffer | 520 |
| Total | 1030 |

Incubate at room temperature for 1 hour and then store at -20 °C.

**5X TAPS-DMF buffer** 50mM TAPS-NaOH pH 8.5, 25mM $MgCl_2$, 50% v/v DMF.

**SPRI-guanidine binding buffer** 4M guanidine thiocyanate, 40mM TRIS, 17.6mM EDTA, pH 8.0. TRIS 1M pH 8 and EDTA 0.5M pH 8 can be added to the 4M guanidine (after the guanidine is solubilized in water – add the proper volume for getting the right final concentration) and then the pH will be very close to 8. Bring the pH to 8 with HCl.

**Sera-Mag Magnetic Beads preparation** Add 1 ml of Sera-Mag Magnetic Beads (Fisher/GE) to a 1.5 ml Eppendorf tube. Place in a magnetic rack. Remove the liquid. Remove the tube from the rack. Add 1 ml of TE and homogenize. Place back in the magnetic rack and remove the liquid. Repeat this step for a total of two TE pH 8.0 washes. Then, add 1 ml of TE pH 8.0. Note: this beads preparation step is required for preparing SPRI-guanidine beads and SPRI-beads.

**SPRI-guanidine beads preparation** Add 10 ml of 5M NaCl to 9 g of PEG 8000 and then add SPRI-guanidine binding buffer (prepared as described above) up to 49 ml. Homogenize during 5 min. Add 1 ml of Sera-Mag Magnetic Beads in TE (prepared as described above) and homogenize. Keep at 4°C.

**SPRI-beads preparation** Add 10 ml of 5M NaCl, 500 µl of 1M TRIS and 100 µl of 0.5M EDTA to 9 g of PEG 8000. Complete the volume to 49 ml with ultra-pure water. Add 1 ml of Sera-Mag Magnetic Beads in TE (prepared as described above) and homogenize. Add 27.5 µl of Tween-20 and homogenize. Keep at 4°C.

**Prepare 2X deamination buffer (as described in Kim et al, 2019, Nature Biotechnology).**

| Component | Final concentration |
|---|---|
| Tris-HCl (pH 8.0) | 100 mM |
| KCl | 50 mM |
| $MgSO_4$ | 5 mM |
| EDTA | 0.2 mM |
| Glycerol | 20 % |
| DTT | 4 mM |
| $ZnCl_2$ | 20 µM |

**PROCEDURE**

**Genomic DNA Isolation**

**1|** Perform genomic DNA isolation with Gentra Puregene Kit (Qiagen), following the manufacturer's instructions.

**CHANGE-seq-BE library preparation**

*Note:* For each sgRNA target site that will be evaluated, 8-10 µg of genomic DNA input for tagmentation is required (8-10 tagmentation reactions), and an additional 8-10 reactions for the control.

*Note:* Tn5-transposome assembly and tagmentation reaction may vary according to the source of Tn5. Optimization might be required for tagmenting the gDNA to ~400 bp.

**2|** Genomic DNA tagmentation. Tn5 reactions are assembled as follows:

| Component | Volume (µl) |
|---|---|
| 5x TAPS-DMF buffer | 20 |
| Tn5 preassembled with oCRL225/oCRL226-MEDS | 40 |
| Genomic DNA (50 ng/µl) | 20 |
| Nuclease-free water | 20 |
| Total | 100 |

Incubate in a thermocycler at 55 °C for 7 minutes.

**3|** Dilute proteinase K 1:1 in water (2.5 µl of proteinase K and 2.5 µl of water) and add 5 µl of the dilution to the tagmented DNA. Incubate at 55 °C for 15 minutes.

**4|** Add 1.8X volumes (189 µl) of SPRI-Guanidine beads to the tagmented DNA, mix thoroughly by pipetting 10 times. Incubate at room temperature for 10 minutes. Place the reaction plate onto a Magnum FLX magnetic rack for 5 minutes. Remove the cleared solution from the reaction plate and discard. Add 200 µl of 80% ethanol, incubate for 30 seconds and remove the supernatant. Repeat this step for a total of two ethanol washes. Remove ethanol completely and let the samples air dry for 3 minutes on the magnetic rack. Remove the plate from the magnetic rack and add 46 µl of TE pH 8.0, and pipette 10 times to mix. Incubate at room temperature for 2 minutes. Place the reaction plate back to the magnetic rack for 1 minute. Transfer the eluted DNA to a new plate, with 23 µl of sample in each well (each tagmentation reaction will be split in two reactions for gap repair Step 6, i.e. if you have 48 samples for tagmentation step 2, you will have 96 samples for gap-repair step 6).

**5|** Run 10 µl (from one reaction only) on a QIAxcel capillary electrophoresis instrument, in a 0.2 ml thin-walled 12-well strip tube with a QIAxcel DNA High Resolution Kit (Qiagen), QX Alignment Marker 15 bp – 10 kb (Qiagen) and QX Size Marker 250 bp – 8 kb (Qiagen), following manufacturer's instruction. The average size of tagmented DNA should be ~400 bp. Quantify by Qubit HS.

**6|** Gap repair. Perform the gap repair reaction as follows:

| **Component** | **Volume (µl)** |
|---|---|
| 2X Kapa HiFi HotStart Uracil+ Ready Mix | 25 |
| Taq DNA ligase | 2 |
| Purified Tagmented DNA (150-250ng) | 23 |
| Total | 50 |

Incubate in a thermocycler at 72°C for 30 minutes.

**7|** Dilute proteinase K 1:1 in water (2.5 µl of proteinase K and 2.5 µl of water) and add 5 µl of the dilution to the gap-repaired DNA. Incubate in a thermocycler at 55°C for 15 minutes.

**8|** Purify the gap repair reactions as previously described in step 4 by adding 1.8X volumes (99 µl) of SPRI-Guanidine beads to the gap repaired-DNA. Elute in 20 µl of TE pH 8.0. Combine every two eluted DNA samples for transferring to a new plate (each sample in the new plate will have 40 µl, meaning that if you have 96 samples for gap-repair n step 6, you will have 48 samples for USER/PNK treatment in step 9).

*Note:* it is crucial to use the SPRI-Guanidine beads in this purification step to completely inactivate and removal of the Kapa HiFi HotStart Uracil+ polymerase, as carryover of this enzyme will remove the 3' overhangs generated in step 9, due to its strong 3'-5' exonuclease activity.

**9|** USER/PNK. Set up the USER/PNK reaction as follows:

| **Component** | **Volume (µl)** |
|---|---|
| T4 DNA Ligase Buffer (10X) | 5 |
| USER Enzyme (1 U/µl) | 3 |
| T4 Polynucleotide Kinase (10 U/µl) | 2 |
| Gap-repaired DNA | 40 |
| Total | 50 |

Incubate in a thermocycler at 37°C for 1 hour.

**10|** Add 1.8X volumes (90 µl) of SPRI-beads to the USER/T4 PNK treated DNA, mix thoroughly by pipetting 10 times. Incubate at room temperature for 10 minutes. Place the reaction plate onto a Magnum FLX magnetic rack for 3 minutes. Remove the cleared solution from the reaction plate and discard. Add 200 µl of 80% ethanol, incubate for 30 seconds and remove the supernatant. Repeat this step for a total of two ethanol washes. Remove ethanol completely and let the samples air dry for 3 minutes on the magnetic rack. Remove the plate from the magnetic rack and add 35 µl of TE pH 8.0, and pipette 10 times to mix. Incubate at room temperature for 2 minutes. Place the reaction plate back to the magnetic rack for 1 minute. Transfer the supernatant to a new plate. Pool and quantify by Qubit dsDNA HS assay.

**11|** Intramolecular circularization. Set up the DNA circularization as follows:

| **Component** | **Volume (µl)** |
|---|---|
| T4 DNA Ligase Buffer (10X) | 10 |
| T4 DNA Ligase (400 U/µl) | 2 |
| USER/PNK treated DNA (500 ng) | *variable* |
| IDTE pH 8.0 | *variable* |
| Total | 100 |

Incubate in a thermocycler at 16 °C for 16 hours.

**12|** Add 1X volumes (100 µl) of SPRI-beads to the circularized DNA, mix thoroughly by pipetting 10 times. Incubate at room temperature for 10 minutes. Place the reaction plate onto a Magnum FLX magnetic rack for 3 minutes. Remove the cleared solution from the reaction plate and discard. Add 200 µl of 80% ethanol, incubate for 30 seconds and remove the supernatant. Repeat this step for a total of two ethanol washes. Remove ethanol completely and let the samples air dry for 3 minutes on the magnetic rack. Remove the plate from the magnetic rack and add 37 µl of TE pH 8.0, and pipette 10 times to mix. Incubate at room temperature for 2 minutes. Place the reaction plate back to the magnetic rack for 1 minute. Transfer the supernatant to a new plate.

**13|** Plasmid-Safe ATP-dependent DNase/Lambda Exo/ExoI/ExoIII treatment:

| **Component** | **Volume (µl)** |
|---|---|
| Ampligase Buffer (10X) | 5 |
| ATP (25 mM) | 2 |
| Plasmid-Safe ATP-Dependent DNase (10 U/µl) | 2 |
| Lambda Exonuclease (5 U/µl) | 2 |
| Exonuclease I (*E. coli*) (20 U/µl) | 1 |
| Exonuclease III (*E. coli*) | 1 |
| Circularized DNA | 37 |
| Total | 50 |

Incubate in a thermocycler at 37 °C for 1 h. Add 5 µl of EDTA 0.5 M, incubate at 70 °C for 30 min, hold at 4 °C.

**14|** Add 1X volumes (50 µl) of SPRI-beads to the circularized DNA, mix thoroughly by pipetting 10 times. Incubate at room temperature for 10 minutes. Place the reaction plate onto a Magnum FLX magnetic rack for 3 minutes. Remove the cleared solution from the reaction plate and discard. Add 200 µl of 80% ethanol, incubate for 30 seconds and remove the supernatant. Repeat this step for a total of two ethanol washes. Remove ethanol completely and let the samples air dry for 3 minutes on the magnetic rack. Remove the plate from the magnetic rack and add 15 µl of TE pH 8.0, and pipette 10 times to mix. Incubate at room temperature for 2 minutes. Place the reaction plate back to the magnetic rack for 1 minute. Transfer the supernatant to a new plate, pool and quantify by Qubit HS. Circularized DNA can be stored at -20°C.

**15|** Quick CIP treatment of circularized exonuclease-treated DNA.

| **Component** | **Volume (µl)** |
|---|---|
| CutSmart Reaction Buffer (10X) | 5 |
| Quick CIP (5 U/µl) | 2 |
| DNA (125 ng) | *X* |
| IDTE pH 8.0 | volume to complete 50 µl |
| Total | 50 |

Incubate at 37 °C for 10 min. Heat inactivate at 80 °C for 20 min. Purify with 1.8X beads, elute in 15 µl.

*Note:* CIP-treated circularized DNA will need to be concentrated in a vacuum centrifuge to 8 ng/µl or higher.

***In vitro* deamination and cleavage of enzymatically purified, circularized gDNA**

**16|** sgRNA dilution and re-fold. Dilute the sgRNA to 9 µM in nuclease-free water and use the follow program on a thermocycler for sgRNA re-fold:

| **Step** | **Temperature** | **Time** | **Cycles** |
|---|---|---|---|
| 1 | 90 °C | 5 min | 1 |
| 2 | 90-25 °C | Ramp rate 2% | |
| Hold | 4°C | | 1 |

**17|** *In vitro* deamination with ABE8e and sgRNA. Setup *in vitro* deamination master-mix:

| Component | Volume (µl) |
|---|---|
| Digenome-seq Buffer (2X) | 25 |
| ABE8e (5 µM) | 3 |
| sgRNA (9 µM) | 5 |
| Total cleavage master-mix | 33 |

Incubate at room temperature for 10 min.

Add circularized DNA (125 ng, volume vary) and nuclease-free water to complete the 50 µl:

| | |
|---|---|
| Cleavage master-mix | 33 |
| Exonuclease Treated DNA (125 ng) | *X* |
| Nuclease-free water | *X* |
| Total | 50 |

Incubate in a thermocycler at 37 °C for 24 h, hold at 4 °C.

**18|** Dilute proteinase K 1:4 in water (1 µl of proteinase K and 4 µl of water) and add 5 µl of the dilution to the *in vitro*-cleaved DNA and incubate in a thermocycler at 37 °C for 15 min.

**19|** Add 1X volumes (55 µl) of SPRI-beads to the *in vitro*-deaminated DNA, mix thoroughly by pipetting 10 times. Incubate at room temperature for 10 minutes. Place the reaction plate onto a Magnum FLX magnetic rack for 3 minutes. Remove the cleared solution from the reaction plate and discard. Add 200 µl of 80% ethanol, incubate for 30 seconds and remove the supernatant. Repeat this step for a total of two ethanol washes. Remove ethanol completely and let the samples air dry for 3 minutes on the magnetic rack. Remove the plate from the magnetic rack and add 40 µl of TE pH 8.0, and pipette 10 times to mix. Incubate at room temperature for 2 minutes. Place the reaction plate back to the magnetic rack for 1 minute. Transfer the supernatant to a new plate.

**20|** Endonuclease V treatment. Setup the Endonuclease V master mix:

| Component | Volume (µl) |
|---|---|
| NEB Buffer 4 (10X) | 5 |
| Endonuclease V (10 U/µl) | 0.25 |
| Nuclease-free water | 4.75 |
| Total master-mix | 10 |

Add 10 µl of endonuclease V master-mix to each eluted DNA sample.

| | |
|---|---|
| Endonuclease V master-mix | 10 |
| ABE8e treated DNA | 40 |
| Total | 50 |

Incubate on a thermocycler at 37 °C for 1 h, then 65 °C for 20 min, hold at 4 °C.

**21|** Add 1.8X volumes (90 µl) of SPRI-beads to the ABE8e/Endonuclease V treated DNA, mix thoroughly by pipetting 10 times. Incubate at room temperature for 10 minutes. Place the reaction plate onto a Magnum FLX magnetic rack for 3 minutes. Remove the cleared solution from the reaction plate and discard. Add 200 µl of 80% ethanol, incubate for 30 seconds and remove the supernatant. Repeat this step for a total of two ethanol washes. Remove ethanol completely and let the samples air dry for 3 minutes on the magnetic rack. Remove the plate from the magnetic rack and add 40 µl of TE pH 8.0, and pipette 10 times to mix. Incubate at room temperature for 2 minutes. Place the reaction plate back to the magnetic rack for 1 minute. Transfer the supernatant to a new plate,

**22|** End repair. Setup the end-repair master mix:

| Component | Volume (µl) |
|---|---|
| NEB Buffer 2 (10X) | 5 |
| Klenow fragment (3'->5' exo-) (5 U/µl) | 1 |
| dNTP | 4 |
| Total end-repair master-mix | 10 |

Add 10 µl end-repair master-mix to each eluted DNA sample.

| | |
|---|---|
| End-repair master-mix | 10 |
| ABE8e treated DNA | 40 |
| Total | 50 |

Incubate on a thermocycler at 37 °C for 30 min, then 75 °C for 20 min, hold at 4 °C.

**23|** Add 1X volumes (50 µl) of SPRI-beads to end-repaired DNA, mix thoroughly by pipetting 10 times. Incubate at room temperature for 10 minutes. Place the reaction plate onto a Magnum FLX magnetic rack for 3 minutes. Remove the cleared solution from the reaction plate and discard. Add 200 µl of 80% ethanol, incubate for 30 seconds and remove the supernatant. Repeat this step for a total of two ethanol washes. Remove ethanol completely and let the samples air dry for 3 minutes on the magnetic rack. Remove the plate from the magnetic rack and add 42 µl of TE pH 8.0, and pipette 10 times to mix. Incubate at room temperature for 2 minutes. Keep the beads.

**24|** A-tailing. Setup the A-tailing master mix (reagents provided with HTP Library Preparation Kit PCR-free (96rxn) (Kapa Biosystems):

| Component | Volume (µl) |
|---|---|
| NEB dA-tailing Reaction Buffer (10X) | 5 |
| Klenow fragment (exo-) | 3 |
| Total A-tailing master-mix | 8 |

Add 8 µl of A-tailing master-mix to each eluted DNA sample with beads.

| | |
|---|---|
| A-tailing master-mix | 8 |
| Cleaved DNA/beads | 42 |
| Total | 50 |

Incubate on a thermocycler at 30 °C for 30 min, hold at 4 °C.

**25|** Add 1.8X volumes (90 µl) of PEG/NaCl SPRI solution (provided with HTP Library Preparation Kit PCR-free (96rxn) (Kapa Biosystems)) to A-tailed DNA, mix thoroughly by pipetting 10 times. Incubate at room temperature for 10 minutes. Place the reaction plate onto a Magnum FLX magnetic rack for 3 minutes. Remove the cleared solution from the reaction plate and discard. Add 200 µl of 80% ethanol, incubate for 30 seconds and remove the supernatant. Repeat this step for a total of two ethanol washes. Remove ethanol completely and let the samples air dry for 3 minutes on the magnetic rack. Remove the plate from the magnetic rack and add 25 µl of TE pH 8.0, and pipette 10 times to mix. Incubate at room temperature for 2 minutes. Keep the beads.

**26|** Adapter ligation. Setup the adapter ligation master-mix (reagents provided with HTP Library Preparation Kit PCR-free (96rxn) (Kapa Biosystems)):

| Component | Volume (µl) |
|---|---|
| NEBNext Ultra II Ligation Master Mix | 30 |
| NEBNext Ligation Enhancer | 1 |
| NEBNext Adapter for Illumina (15 µM) | 2.5 |

| | |
|---|---|
| Total master-mix | 33.5 |

Add 25 µl of adapter ligation master-mix to each A-tailed DNA sample with beads.

| | |
|---|---|
| Adapter ligation master-mix | 33.5 |
| A-tailed DNA/beads | 60 |
| Total | 93.5 |

Incubate on a thermocycler at 20 °C for 15 min (do not heat the lid), hold at 4 °C.

*Note*: Prepare single-use aliquots of NEB adapters to avoid adapter dimer formation due to freeze–thaw hydrolysis of the 3′ T′.

**27|** Add 1X volumes (50 µl) of PEG/NaCl SPRI solution (provided with HTP Library Preparation Kit PCR-free (96rxn) (Kapa Biosystems)) to the adapter-ligated DNA and purify DNA as described in step 20. Elute in 47 µl of TE pH 8.0 and keep the beads.

**28|** USER enzyme. Add 3 µl of USER enzyme, provided with NEBNext® Multiplex Oligos for Illumina® (Dual Index Primers Set 1) to the adapter ligated DNA with beads. Incubate at 37 °C for 15 min.

**29|** Add 0.7X volumes (35 µl) of PEG/NaCl SPRI solution (provided with HTP Library Preparation Kit PCR-free (96rxn) (Kapa Biosystems)) to the USER Enzyme treated DNA and purify as previously described in step 20. Elute in 20 µl of TE pH 8.0. Transfer the supernatant to a new semi-skirted PCR plate and quantify by Qubit dsDNA HS assay and proper Qubit assay tubes (usually about 2-5 ng/µl).

**30|** PCR. Setup a PCR master-mix for adding dual-index barcodes:

| **Component** | **Volume (µl)** | **Final concentration** |
|---|---|---|
| Nuclease-free water | 5 | |
| 2X Kapa U+ HiFi HotStart Ready Mix | 25 | 1X |
| Total master-mix | 30 | |
| Diluted PCR master-mix | 30 | |
| NEBNext i5 Primer (10 µM) | 5 | 1 µM |
| NEBNext i7 Primer (10 µM) | 5 | 1 µM |
| Total PCR mix | 45 | |

**31|** Add 40 µl of PCR master-mix to each sample of purified, USER enzyme treated DNA (~20ng).

| **Component** | **Volume (µl)** | **Final concentration** |
|---|---|---|
| PCR mix | 40 | |
| USER enzyme treated DNA (~10-20ng) | 10 | ~0.4 ng/µl |
| Total | 50 | |

**32|** Perform the PCR using the following thermocycling conditions:

| **Step** | **Temperature** | **Time** | **Cycles** |
|---|---|---|---|
| Denaturation | 98 °C | 45 s | 1 |
| Denaturation | 98 °C | 15 s | 20 |
| Annealing | 65 °C | 30 s | 20 |
| Extension | 72 °C | 30 s | 20 |
| Extension | 72 °C | 1 min | 1 |

| | | |
|---|---|---|
| Hold | 4°C | 1 |

**33|** Purification Add 0.7X volumes (35 µl) of SPRI-beads to the PCR and purify as previously described in step 18. Elute in 30 µl of TE pH 8.0. Transfer the supernatant to a new semi-skirted PCR plate and run 3 µl in QIAxcel.

**34|** Make 1:10 serial dilutions of 50 µl from $10^{-1}$ to $10^{-5}$ dilution of each sample from the library (PCR), starting with 5 µl of DNA and 45 µl of nuclease-free TE pH 8.0, and mix well.

**35|** Assemble qPCR master-mix solution as follows:

| Component | Volume (µl) | Final Concentration |
|---|---|---|
| KAPA SYBR FAST qPCR Master Mix (2X) + Primer Premix (10X) | 12 | 1X |
| Nuclease-free water | 4 | |
| Total qPCR mix | 16 | |

**36|** Assay 2 different dilution factors (4 µl) for each sample ($10^{-4}$, and $10^{-5}$ from the library) in duplicate (in an appropriate 96-well plate). A standard curve (provided with Kapa Library Quantification Kit) and a non-template control (NTC) are required. Add 4 µl of each standard in duplicate, and nuclease-free water in the NTC. Add 16 µl of qPCR master-mix to each sample.

| Component | Volume (µl) | Final concentration |
|---|---|---|
| qPCR mix | 16 | |
| Sample (add nuclease-free water into the NTC well) | 4 | *variable* |
| Total | 20 | |

**37|** Seal the plate and spin down.

**38|** Run qPCR in appropriate thermocycler with the following program:

| Cycling step | Temperature | Time | Cycles |
|---|---|---|---|
| Initial denaturation | 95 °C | 5 min | 1 |
| Denaturation | 95 °C | 30 s | 35 |
| Annealing/extension/data acquisition | 60 °C | 45 sec | 35 |
| Melt curve analysis | 60-95 °C | | |

**39|** Add the appropriate DNA copies for each standard when setting up the qPCR plate in the qPCR program, as follows:

| Standard | dsDNA molecules/µl |
|---|---|
| Standard 1 | $1.2x10^{7}$ |
| Standard 2 | $1.2x10^{6}$ |
| Standard 3 | $1.2x10^{5}$ |
| Standard 4 | $1.2x10^{4}$ |
| Standard 5 | $1.2x10^{3}$ |
| Standard 6 | $1.2x10^{2}$ |

**40|** Analyze qPCR results. Multiply the average of duplicate values by the dilution factor and by the five-fold dilution factor of the qPCR reaction, as follows: Total copies/µl = # * dilution factor.

**41|** Pool library for NextSeq. Pool all the samples in one library at equimolar concentrations. 1X pooled library should be in a total volume of 5 µl, ~ $8 x 10^{9}$ molecules.

**42|** Denature the pooled library (~ 8 x $10^9$ molecules) by adding 5 µl of NaOH 0.2N and incubate at room temperature for 5 min. Then, 5 µl of 200 mM Tris-HCl pH 7.0 and 985 µl of NextSeq buffer (supplied with NextSeq® Reagent Kit).

**43|** Prepare the Phix control V3 (PhiX Control V3 KIT) as follows: mix 2 µl of 10 nM PhiX control with 3 µl of Tris-HCl 10 mM + 0.1% Tween-20, denature with 5 µl of NaOH 0.2N and incubate at room temperature for 5 min. Then, add 5 µl of 200 mM Tris-HCl pH 7.0 and 985 µl of NextSeq buffer (supplied with NextSeq® Reagent Kit).

**44|** Take 120 µl of denatured library, and add 15 µl of denatured Phix. Complete the volume to 1.3 ml with NextSeq buffer.

**45|** Clean the Flow Cell (supplied with NextSeq® Reagent Kit) with alcohol pad and lens wipe.

**46|** Load and sequence library using a NextSeq 300-cycles Mid or High output kit according to manufacturer's instructions using NextSeq 550 system. Sequencing is performed with 151 bp paired-end reads and 8 bp dual-index reads. At least 20M sequencing reads are required per sample.

**47|** After sequencing, copy the demultiplexed output FASTQ files to a location accessible to CIRCLE-seq/CHANGE-seq analysis pipeline.

**Supplementary Protocol References**

1. Lazzarotto, C. R. *et al.* CHANGE-seq reveals genetic and epigenetic effects on CRISPR-Cas9 genome-wide activity. *Nat. Biotechnol.* **38**, 1317–1327 (2020).
2. Newby, G. A. *et al.* Base editing of haematopoietic stem cells rescues sickle cell disease in mice. *Nature* **595**, 295–302 (2021).
